# Prisoner of War dynamics explains the time-dependent pattern of substitution rates in viruses

**DOI:** 10.1101/2021.02.09.430479

**Authors:** Mahan Ghafari, Peter Simmonds, Oliver G Pybus, Aris Katzourakis

## Abstract

Molecular clock dating is widely used to estimate timescales of phylogenetic histories and to infer rates at which species evolve. One of the major challenges with inferring rates of molecular evolution is the observation of a strong correlation between estimated rates and the timeframe of their measurements. Recent empirical analysis of virus evolutionary rates suggest that a power-law rate decay best explains the time-dependent pattern of substitution rates and that the same pattern is observed regardless of virus type (*e.g.* groups I-VII in the Baltimore classification). However there exists no explanation for this trend based on molecular evolutionary mechanisms. We provide a simple predictive mechanistic model of the time-dependent rate phenomenon, incorporating saturation and host constraints on the evolution of some sites. Our model recapitulates the ubiquitous power-law rate decay with a slope of −0.65 (95% HPD: −0.72, −0.52) and can satisfactorily account for the variation in inferred molecular evolutionary rates over a wide range of timeframes. We show that once the saturation of sites starts - typically after hundreds of years in RNA viruses and thousands of years in DNA viruses - standard substitution models fail to correctly estimate divergence times among species, while our model successfully re-creates the observed pattern of rate decay. We apply our model to re-date the diversification of genotypes of hepatitis C virus (HCV) to 396,000 (95% HPD: 326,000 - 425,000) years before present, a time preceding the dispersal of modern humans out of Africa, and also showed that the most recent common ancestor of sarbecoviruses dates back to 23,500 (95% HPD: 21,100 - 25,300) years ago, nearly thirty times older than previous estimates. This not only creates a radical new perspective for our understanding the origins of HCV but also suggests a substantial revision of evolutionary timescales of other viruses can be similarly achieved.

## Main

The timescale over which viruses evolve and how this process is connected to host adaptation has been an area of considerable research and methodological progress in recent decades. Mammalian RNA viruses, in particular, exhibit extraordinarily rapid genomic change^1–3^ and analyses of their genetic variation has enabled detailed reconstruction of the emergence of viruses such as HIV-1^4^, hepatitis C virus^5^ and influenza A virus^6,7^. RNA viruses display evolutionary change over short-timescales (weeks to months) and can re-model a substantial part of their genomes following a host switch^8–13^. Well characterised examples in both RNA and DNA viruses include the emergence of HIV-1 in humans from a chimpanzee reservoir^4,14,15^, and the adaptation of myxomatosis in rabbits^16^.

These rapid rates of virus sequence change stand in striking contrast with evidence for extreme conservation of virus genome sequences over longer periods of evolution and at higher taxonomic levels^17,18^. Inferred short term rates of virus sequence change should create completely unrecognisable genome sequences if they were extrapolated over thousands or even hundreds of years, yet endogenous viral elements (EVEs) that integrated into host genomes throughout mammalian evolution are recognisably similar to contemporary genera and families of *Bornaviridae*, *Parvoviridae* and *Circoviridae* amongst many examples^19–24^. This observation is complemented by increasing evidence from studies of virus / host co-evolution^25–27^, and more recently from analysis of viruses recovered from ancient DNA in archaeological remains^28–32^, that together indicate a remarkable degree of conservatism in viral genome sequences and their inter-relationships at genus and family levels. This conundrum has been attributed to the time-dependent rate phenomenon (TDRP), which is the observation that apparent rates of evolution are dependent on timescale of measurement. The TDRP has been explained by processes such as sequence site saturation, short-sighted within-host evolution, short-term changes in selection pressure and potential errors in estimation of short-term substitution rates^17,18,33–35^. Empirically, substitution rates show a striking linear relationship between log-transformed rates and timescale of measurement, across RNA and DNA viruses, despite the large differences among viruses in their initial short term substitution rates^33^. The gradient of regressions of observation time to estimated evolutionary rates is consistently around −0.65, for all virus groups for which long term substitution rates can be calculated or inferred, implying a common underlying evolutionary process.

We recently developed a model of virus evolution in which the primary driver of sequence change over long evolutionary time-scales was host adaptation, in which virus sequence change is severely curtailed by stringent fitness constraints^36^. Viruses exist within a tightly-constraining host niche to which they rapidly adapt; paradoxically, their high mutation rates, large population sizes and consequent ability to adapt rapidly serve to restrict their long-term diversification and sustained sequence change, rendering them evolutionary “Prisoners of War” (PoW). Ultimately over longer timescales, rates of viral evolution will be bounded by the rate of evolution of their hosts ^36^.

In the current study, we develop the PoW model to explain the longer-term evolutionary trajectories of viruses and the time-dependence of their inferred substitution rates. The model accounts for genetic saturation of rapidly-evolving sites, and host constraints on site evolution, with the proportion of fast- to slow-evolving sites being exponentially distributed. These sites will saturate chronologically from the fastest evolving to those that evolve epistatically, to those that evolve at the host substitution rate. This model can reproduce effectively the empirically observed TDRP patterns and the inflection points where time-dependent rate changes become manifest. We demonstrate that the model predictions are robust to intrinsic and marked differences in substitution rates among different virus groups or assumptions about the relative proportion of sites evolving at different rates.

### Power-law rate decay due to site-saturation

First, we show how a time-dependent rate effect emerges when estimating the sequence divergence using a standard evolutionary model. Suppose that a sequence has diverged from its ancestor *t* generations ago under a constant and uniform substitution rate, *μ*. The proportion of pairwise differences between the sequence and its ancestor, *p*(*t*), reaches its maximum value, hereafter called the saturation frequency, *α*, at time *t*\* ≈ *α* / *μ*(see **Equation S1**). As the ancestral and derived sequence continue to evolve beyond the saturation point, *t* *, their observed proportion of pairwise differences, 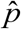, is effectively unchanged. Thus, using any conventional substitution model, the inferred genetic distance, 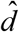, remains constant and the estimated substitution rate, 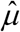, follows a power-law drop as the observed divergence time, 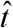, increases, i.e. 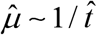, with slope −1 on a log-log graph (see **Figure S1 a**) – this is assuming the substitution model correctly infers the saturation frequency, i.e. *α_M_* ≥ *α*, where *α_M_* is the expected saturation frequency set by the model (see **Equation S2**).

### Saturation under rate heterogeneity

In the presence of among-site rate heterogeneity, a fraction of sites may evolve at a rate that is much slower (or faster) than some other sites. Under such circumstances, if we apply a measure of genetic distance based on a constant and homogeneous substitution rate per site, the substitution model may reliably recover the expected (mean) rate up to and before the fastest-evolving sites reach saturation, beyond which point the time-dependent pattern of rate decay emerges while the remaining sites continue to accumulate changes until the slowest-evolving sites also reach saturation (i.e. the point at which the entire sequence space has been fully explored) and a power-law rate decay with slope −1 emerges (see **Figure S1 b**).

Although our focus so far has been on the saturation of observed pairwise differences and how it can create a time-dependent rate effect, the same holds true when tracking the evolutionary changes of a large number of sequences through time. Using a standard Jukes-Cantor substitution model on a set of simulated sequences (see Methods section), both in the absence and presence of rate heterogeneity, we can recreate similar patterns of time-dependent rate decay and show that, over longer timescales, i.e. when the divergence time between two populations is much longer than the typical coalescent times (2*N_e_*) of a neutral population of size *N_e_*, the variation in inferred substitution rates is dominated by the saturation along the longest (internal) branch connecting the two populations (**Figure S2**). We also find that, over short timescales, systematic under-estimation of the Time to the Most Recent Common Ancestor (TMRCA) results in inflated substitution rate estimates (**Equation S4**).

### Saturation under the Prisoner of War model

The PoW model of virus evolution is based on the principle that site saturation over long timescales dominates the time-dependency of virus substitution rates. Building upon a collection of virus evolutionary rate estimates from >130 publications^33^, we use 389 nucleotide substitution rate estimates across six major viral groups to find the line of best fit between our predicted time-dependent substitution rate (the PoW model) and the evolutionary rate estimates for each viral group (data) using the geometric least squares method^37^. We then show that the PoW model captures all of the important properties of the TDRP and its variation among different groups of viruses.

The model categorises into *M* discrete rate classes that are equally spaced on a log-scale, with a common ratio Δ*M* between consecutive rate groups, ranging from those evolving the fastest, at rate *μ*_max_ per site per year (SSY), to the ones evolving at the host substitution rate, *μ*_min_. The fraction of sites, *m_i_*, in each rate group *i*, with the corresponding substitution rate, *μ*_i_, is exponentially distributed, *m_i_* = *Ce^λi^*, where *C* is the normalisation factor, i.e. 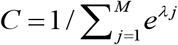, and the exponent coefficient, *λ*, sets the tendency of sites to be either mostly slowly ( *λ* < 0) or rapidly ( *λ* > 0) evolving (**Figure 1 a**). Assuming a fixed incremental difference between any two consecutive rate groups, i.e. *μ*_i+1_ = Δ*M μ_i_*, the substitution rate at the fastest-evolving sites is determined by the total number of groups, *M*, which, in turn, sets the inflection point for when the time-dependent rate decay emerges. Once the fastest-evolving sites diverge to saturation, other rate groups that evolve more slowly (e.g. via epistatic and compensatory substitutions), saturate sequentially as the timespan of rate measurement, *t*, increases. This chronological saturation effect continues until the inferred rate decays to the host substitution rate, *μ*_min_ (**Figure 1 b**). Therefore, the time-dependent rate curve, according to the PoW model is given by

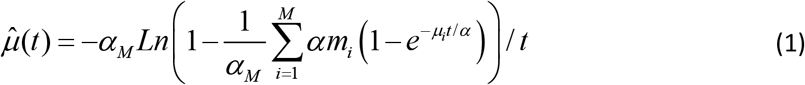

where the observed genetic distance between the derived and ancestral sequences is assumed to increase with the timespan of rate measurement, *t*. While over short timescales, i.e. *t* ≪ 1/ *μ*_max_, several methodological (e.g. internal node calibration errors) and biological (e.g. purifying selection) artefacts may inflate the substitution rate estimates in viruses (i.e. such that 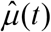 underestimates the inferred substitution rates), over longer time-scales (i.e. after a few years) the rate estimates are expected to converge to the mean substitution rate^38,39^, 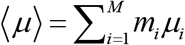.

**Figure 1:**
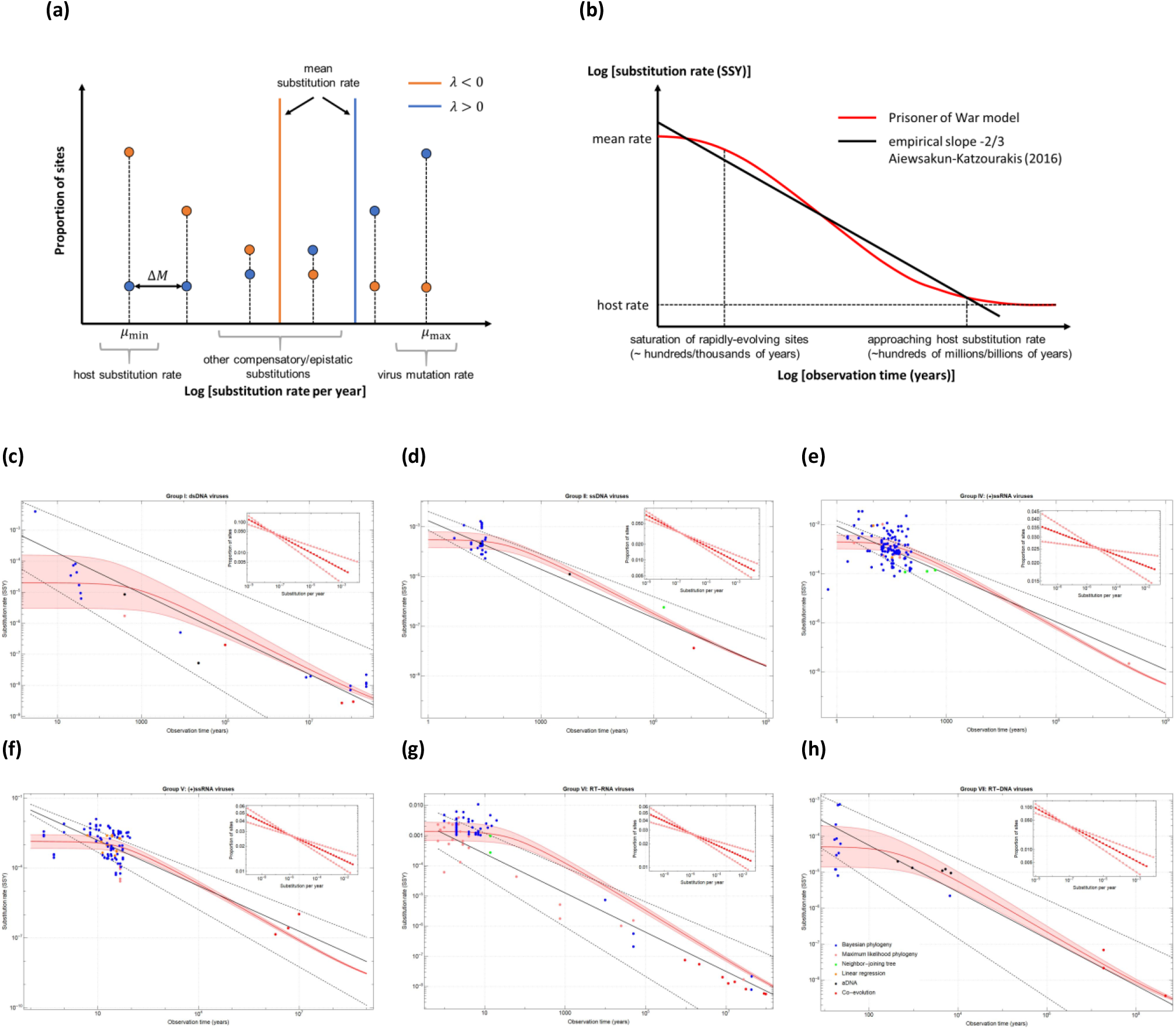
**(a)** Distribution of the fraction of sites per rate group according to the PoW model. A fraction of sites, *m_i_*, belonging to rate group *i* (evolving at rate *μ_i_*), is an exponentially distributed number with parameter *λ*. The rate groups are equally spaced on a log-scale from the slowest (with fixed value), *μ*_min_ = 10^−9^, to the fastest (variable), *μ*_max_, with a common ratio, Δ*M*. The finer the value of Δ*M* gets, the more accurate the predicted values become. However, typically, less than 50 rate groups (i.e. *M* < 50) is sufficient for all model predictions. For a fixed Δ*M*, the exponent coefficient, *λ*, together with the number of rate group, *M*, are the two free parameters of the PoW model which set the mean, 〈*μ*〉, and maximum rates, *μ*_max_, for any given data set. (**b**) Schematic plot of the time-dependent rate dynamics according to the PoW model (red) and the empirical observation made by Aiewsakun and Katzourakis^33^ (black). (**c**)-(**h**) Estimated time-dependent rate curves for each viral group according to the PoW model. A total of 389 viral rate estimates (coloured circles representing various phylogenetic methods used for estimating rates) was collected from more than 130 publications, 23 estimates for group I, 32 for group II, 123 for group IV, 106 for group V, 85 for group VI, and 20 for group VII. The inset shows the distribution of rate groups in each virus group. The red line shows the best fit and shaded area the 95% confidence interval ( Δ*M* = 1.58 and *α*_M_ = *α* = 3 / 4).

Upon the saturation of the fastest-evolving sites (i.e. the inflection point), 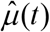 follows the empirically observed power-law decay with the slope −0.65 (95% HPD = −0.72, −0.57) across all viral groups (**Figure 1c-h**). Finding the exact point at which the rate decay pattern emerges likely depends on the choice of substitution model and varies between different virus groups. For instance, a previous study^40^ has shown that substitution models with Gamma-distributed rate heterogeneities across sites may perform better at estimating the mean substitution rate over longer timescales, thereby delaying the emergence of the inflection point in the power-law rate decay, compared to models with a strict molecular clock.

We find that the mean substitution rates and fastest-evolving rate groups in double-stranded DNA viruses (dsDNA) is the lowest among all viral groups and that, together with reverse-transcribing DNA (RT-DNA) and single-stranded DNA viruses (ssDNA), dsDNA viruses have typically 1-2 orders of magnitude slower rates than RNA viruses (see **Table 1**). Conversely, the estimated mean and fastest rates are very similar among the groups of positive-strand RNA viruses (+ssRNA), negative-strand RNA viruses (-ssRNA), and RNA retroviruses (RT-RNA). The estimated substitution rates for rapidly evolving sites in RNA viruses, i.e. *μ*_max_ ~ 10^−2^ SSY, resembles their inferred mutation rates^41^ which may also result in their rapid saturation after a few decades^42^.

**Table 1:**
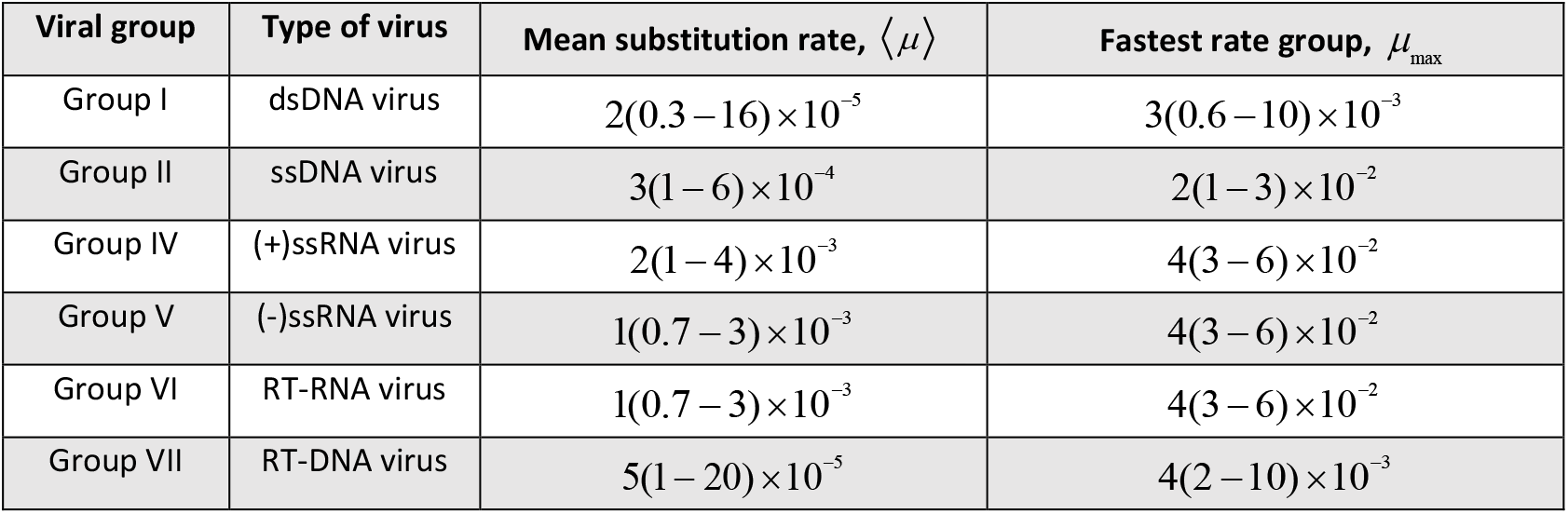
Estimated mean and maximum substitution rate SSY according to the PoW model across 6 viral groups. Parentheses correspond to 95% confidence intervals.

| **Viral group** | **Type of virus** | **Mean substitution rate, $\langle \mu \rangle$** | **Fastest rate group, $\mu_{\max}$** |
| --- | --- | --- | --- |
| Group I | dsDNA virus | $2(0.3 - 16) \times 10^{-5}$ | $3(0.6 - 10) \times 10^{-3}$ |
| Group II | ssDNA virus | $3(1 - 6) \times 10^{-4}$ | $2(1 - 3) \times 10^{-2}$ |
| Group IV | (+)ssRNA virus | $2(1 - 4) \times 10^{-3}$ | $4(3 - 6) \times 10^{-2}$ |
| Group V | (-)ssRNA virus | $1(0.7 - 3) \times 10^{-3}$ | $4(3 - 6) \times 10^{-2}$ |
| Group VI | RT-RNA virus | $1(0.7 - 3) \times 10^{-3}$ | $4(3 - 6) \times 10^{-2}$ |
| Group VII | RT-DNA virus | $5(1 - 20) \times 10^{-5}$ | $4(2 - 10) \times 10^{-3}$ |

To ensure that our model predictions are not biased towards a particular virus family with more evolutionary rate estimates (i.e. more data points to fit to the PoW model in **Equation 1**), we remove all the short-term rate estimates (< 100 years) within each viral group except for one virus family or genus to re-calibrate the mean substitution rates (see **Table S1**). We find that despite the broad range of evolutionary rate estimates across all viral groups, the estimated parameters of the PoW model are robust to such changes and are not an artifact of systematic biases in selecting rate estimates from a particular virus family. We note that, in group VI, the stark difference in evolutionary rates between *Lentivirus* and *Deltaretrovirus* families over short timescales results in a noticeably different pattern of rate decay (see **Figure S3 e**) and the long-term rates are more aligned with the predictions based on the *Deltaretrovirus* re-calibration. The predicted mean and maximum substitution rate of *Lentivirus* families are 1-2 orders of magnitude higher than the *Deltaretrovirus* family. The latter evolves at rates similar to RT-DNA viruses. We also see that a larger fraction of sites in DNA viruses tend to evolve at rates closer to the host substitution rates (i.e. have a sharper negative gradient, *λ*) compared to RNA viruses, which largely have an equal fraction of sites across all rate groups. It is worth noting that all rate groups, including the ones with lowest proportion of sites, *m*_min_, have representative proportions across the genome, i.e. *m*_min_ ≫ 10^−4^ > 1/ *L*, where *L* is a typical genome size of an RNA virus. We also carried out a similar sensitivity analysis at the level of virus genera which further confirms that the rate curves predicted by the PoW model are still accurate at this level and are not an artefact of measured rates at the level of Baltimore groups (see **Figure S4**).

To illustrate the radical effect of applying the PoW model to virus evolutionary timescales, we analysed an alignment of complete hepatitis C virus (HCV) genome sequences that represent its component genotypes and subtypes (**Figure 2**). Using the trajectory of the PoW-transformed evolutionary rate decay for Group IV and the expected (short-term) substitution rate of 1.2 x 10^−3^ substitutions/site/year for HCV (**Figure 2 a, b**), the model demonstrates a clear separation of timescales for the diversification of variants within genotypes ( ~ 50 - 500 years), among subtypes ( ~ 1, 000 - 20, 000 years), and among genotypes ( ~ 40, 000 - 200, 000 years) with an estimated root age for HCV of 396,000 (95% HDP: 326,000 – 425,000) years before present (BP). While the predicted divergence times for within-genotype variants using the PoW model is similar to those obtained using a standard substitution model, the latter models estimate the root age of HCV to be only 970 (95% HDP: 850 – 1100) years BP with no clear separation of timescales for among-genotype diversifications (**Figure 2 c**). These results contrast with estimates of 500 – 2,000 years of genotype diversification by simple extrapolation from short term rates^43^, while among-subtype divergence times of 1,000 – 20,000 years are up to 50 times higher than the 300 – 500 years estimated in previous molecular epidemiological analysis^44–46^. The revised, very early evolutionary origin of HCV genotypes (326,000years, 425,000 years 95% HPD) predicted by our model is striking. While these early dates still fit currently proposed hypotheses for multiple and potentially relatively recent zoonotic sources of HCV in humans, associated with different genotypes^47,48^, the existence of a common ancestor of HCV before human migration of Africa (150,000BP) support alternative scenarios where HCV diversified within anatomically modern humans. HCV genotypes may have arisen from geographical separation in Africa (genotypes 1, 2, 4, 5, 7) and migrational separation of human populations migrating out of Africa into Asia (genotypes 3, 6 and 8).

**Figure 2:**
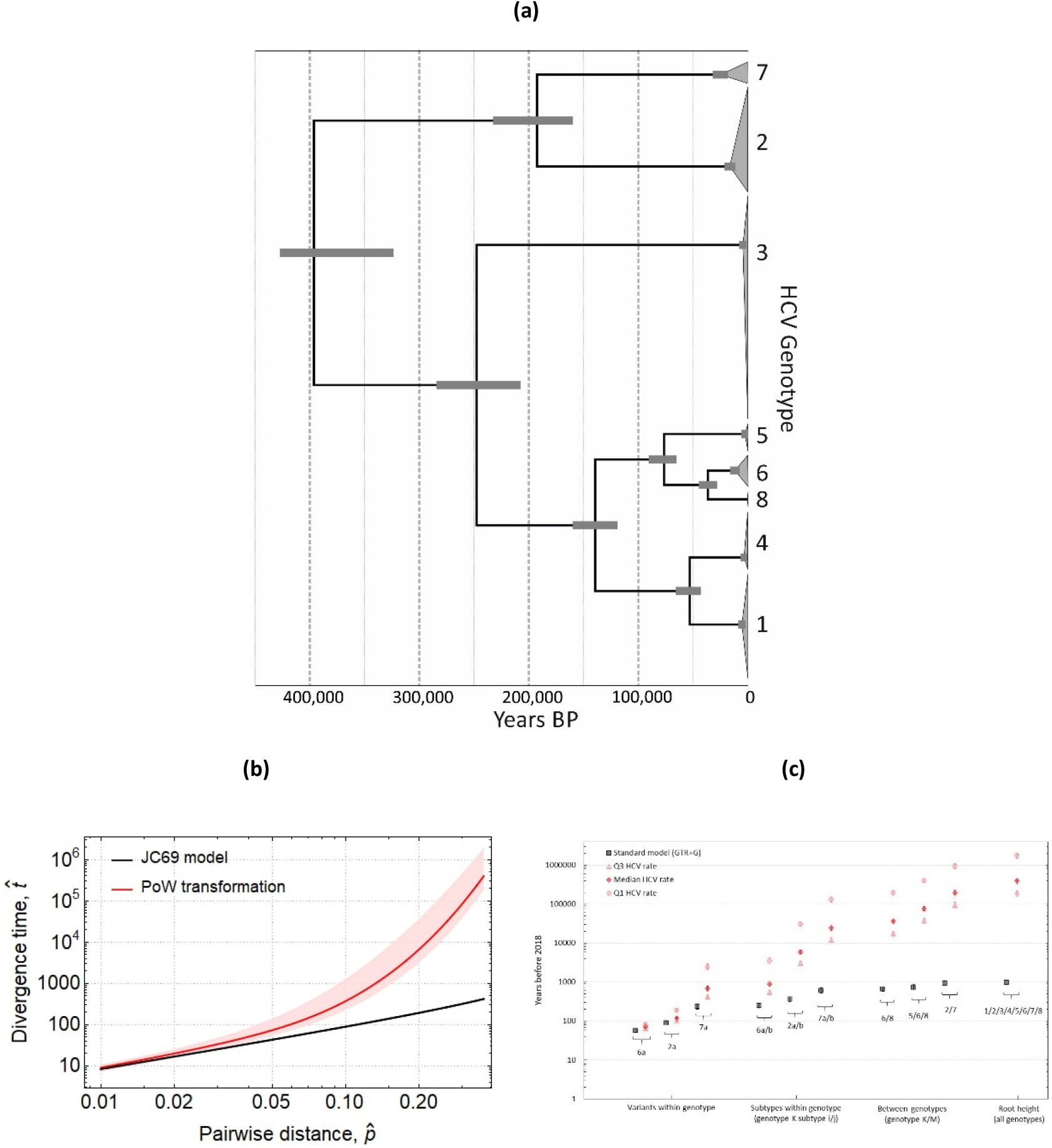
**(a)** The PoW-transformed time-calibrated phylogeny represents a maximum clade credibility tree inferred for HCV (including all its 8 genotypes). An ultrametric tree was first estimated in the Bayesian phylogenetic framework under a strict clock assumption and a JC69 substitution model assuming a fixed substitution rate equal to 1 (branch lengths are in units of substitutions per site) using BEAST software platform (v.1.10)^52^. Then the branch lengths (along with their corresponding 95% HPD) are rescaled according to the PoW transformation. The two insets show magnified parts of the tree where genotypes 5, 6, 7, and 8 along with their subtypes (i.e. a and b) and nearest within-genotype variants (e.g. MH940291/-/2015 is a variant within genotype 7a) are highlighted. **(b)** Compares the estimated divergence time for a pair of HCV sequences as a function of their inferred genetic distance (ranging from 1% to 37%) using a standard JC69 substitution model with an estimated distance 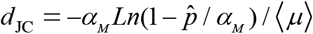 and a PoW-transformed measure of distance. The expected (short-term) HCV substitution rate (taken out of 9 estimates from the literature^33^) is 〈*μ*〉 = 1.2(IQR: 1.1-1.4)×10^−3^ SSY and *μ*_max_ is assumed to be the same as the one inferred for Group IV (see **Table 1**). (**c**) Shows the estimated divergence times within and between various HCV genotypes and subtypes using a general time-reversible substitution model with a four-bin gamma rate distribution (GTR+G) and PoW-transformed tree. Error bars represent the 95%HDP for the inferred maximum clade credibility tree.

We also carried out a similar analysis to investigate the origins of the SARS-CoV-2 sarbecovirus lineage (**Figure 3**). While the PoW-transformed phylogeny recovers the previous estimates for SARS-CoV-1 and SARS-CoV-2 diversification from their most closely related bat virus over short timescales (i.e. less than hundreds of years BP), it extends the root age back to 23,000 (21,000 - 25,000) years BP, nearly 20 times older than previous estimates^49^. Our results indicate that humanity may have been exposed to these viruses since the Paleolithic if they had come into contact with their natural hosts. Also, our date estimates of the origin of the sarbecovirus lineage are in remarkable concordance with signatures of selection on human genomic datasets that indicate an arms race with corona-like viruses dating back 25,000 years^50^, providing an external comparator for our methodology.

**Figure 3:**
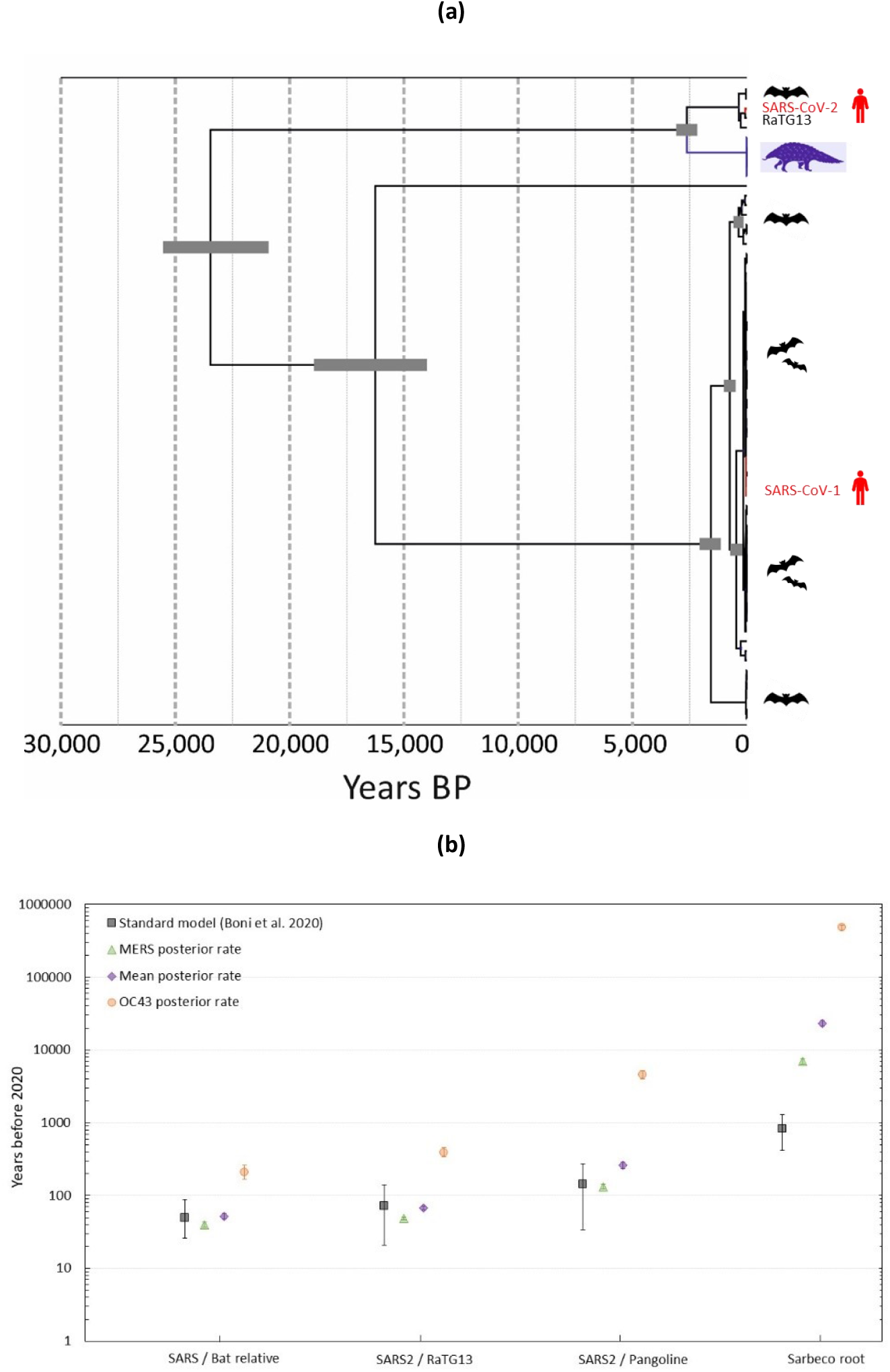
**(a)** The PoW-transformed time-calibrated phylogeny represents a maximum clade credibility tree inferred for the SARS-CoV-2 sarbecovirus lineage based on the non-recombinant alignment 3 (NRA3)^49^ which includes SARS-CoV-1 and SARS-CoV-2 viruses in humans among other closely related bat and pangolin viruses. An ultrametric tree was first estimated in the Bayesian phylogenetic framework under a strict clock assumption and a JC69 substitution model assuming a fixed substitution rate equal to 1 (branch lengths are in units of substitutions per site) using BEAST software platform (v.1.10)^52^. Then the branch lengths (along with their corresponding 95% HPD) are rescaled according to the PoW transformation. The two insets show magnified parts of the tree where SARS-CoV-1 (blue) and SARS-CoV-2 (red) are located. **(b)** Shows the divergence time estimates for SARS-CoV-2 and 2002-2003 SARS-CoV from their most closely related viruses according to a standard substitution model^49^ (black) and the PoW-transformed phylogeny using the consensus posterior rate centred around *μ* = 5.5 ×10^−4^ SSY (purple), HCoV-OC43 rate prior (orange), 2.4 ×10^−4^ SSY, and MERS-CoV rate prior (green), 7.8 ×10^−4^ SSY (see Extended data figure 3 in ref^49^).

The PoW model creates an over-arching evolutionary framework that can reconcile, and incorporate timescales derived from co-evolutionary and ancient DNA studies. Further radical re-evaluations of timescales of other RNA and DNA viruses using this approach will provide new insights into their origins and evolutionary dynamics^27,51^. Application of the PoW will place ancestors of more divergent virus sequences far further back into the past than conventional reconstructions. The good fit between modelled and observed substitution rates and the gradient of rate decay over time were based on a minimal number of assumptions about mutational fitness effects, proportion of sites evolving at a particular rate, and robust to substantial differences in substitution rates across different viral groups. By finding the short-term substitution rate (the flat part of the modelled rate decay) and the value of the fastest-evolving rate group (which sets the inflection point of the curve), the PoW model can reconstruct corrected substitution rates for virus genotypes with increasingly divergent nucleotide sequences.

## Methods

### Time-dependent rate decay under a uniform substitution rate

Suppose that a sequence has diverged from its ancestor *t* generations ago under a substitution rate *μ* that is constant over time and equal across all sites. In this case, the proportion of pairwise differences, *p*(*t*), is given by^53^

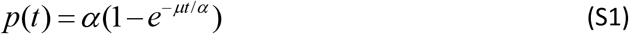

such that *α* is the maximum proportion of pairwise differences and is determined by 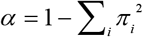 where *π_i_* is the base frequency of the *i* th nucleotide or amino acid. Assuming that *d* is the ‘true’ genetic distance between a pair of homologous sequences, i.e. *d* = *μt*, we can estimate the observed genetic distance, 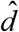, with an observed proportion of pairwise differences, 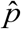, using the Felseinstein’s 1981 substitution model^54^

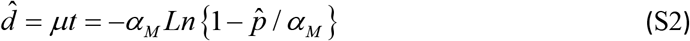

where *α_M_* is the expected saturation frequency set by the model. If the model correctly estimates the saturation frequency, i.e. *α_M_* = *α*, **Equation S2** accurately predicts the true genetic distance, i.e. 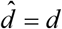, as long as the divergence time *t* ≪ *α* / *μ*. As *p*(*t*) approaches saturation frequency at *t* ≈ *α*/ *μ*, the observed proportion of pairwise differences will be bound by the number of evolving sites. For instance, if the saturation frequency is *α* = 3 / 4, i.e. a standard Jukes-Cantor substitution model, to distinguish between an observed pairwise difference of 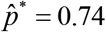 and 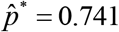 would require a sample of approximately one thousand evolving site at rate *μ*. Thus, for any small sample size, the estimated distance, 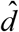, will remain effectively unchanged for *t* ≳ *α*/*μ*. In other words, if the pair have evolved beyond their saturation point, for a given divergence time, 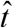, the inferred rate follows a power-law rate drop with a slope −1 on a log-log plot, i.e. 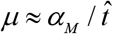(see grey curve in **Figure S1 a**). The same power-law bias can be observed when estimating the divergence time, 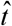, assuming there is prior knowledge on the inferred substitution rate, 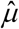.

In the presence of purifying selection and/or amino acid and nucleotide biases, the substitution model in **Equation S2** may incorrectly assume a maximum proportion of differences, *α_M_*, that is higher than the true value, i.e. *α_M_* > *α*. For instance, if we apply a Jukes-Cantor measure of nucleotide distance to a pair of sequences with a particular site preference that equally favours only two (out of the four) nucleotides, i.e. *α* = 1/ 2, the true proportion of differences reaches saturation at earlier divergence times than what the selected substitution model would predict. As a result, similar to the previous scenario, the estimated rate drops as a power-law with slope −1 (see orange curve in **Figure S1 a**) after hitting the saturation point, i.e. 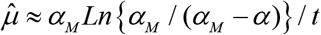.

The reverse side of this is when *α_M_* < *α* in which case the model underestimates the true saturation level. In this case, the observed proportion of pairwise differences passes the substitution model’s expected point, i.e. 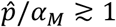, at which point the estimated substitution rate, 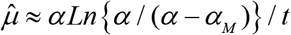, goes to infinity (see blue curve in **Figure S1 a**).

### Time-dependent rate decay in the presence of rate heterogeneity

In the previous scenario, **Equation S1**, we assumed there is no rate heterogeneity across sites. However, if the actual evolutionary process involves *M* group of sites with each group *i* evolving at rate *μ_i_* and occupying a fraction *m_i_* of sites, the proportion of pairwise differences would be given by

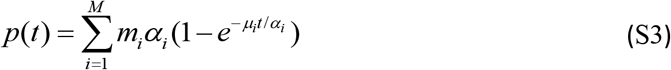

such that ∑_*i*_ *m_i_* = 1 and the expected substitution rate 〈*μ*〉 = ∑*_i_ m_i_ μ_i_*. If we apply a measure of distance based on **Equation S2** to an evolutionary process with rate heterogeneity, **Equation S3**, the model can reliably infer the expected substitution rate up to when the first group of sites with the fastest substitution rate, *μ*_max_, reach saturation at 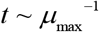at which point the pattern of time-dependent rate decay emerges until it plateaus at a rate corresponding to the slowest-evolving sites, *μ*_min_. Once all the rate groups reach saturation at time 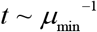, the power-law rate decay with slope −1 emerges.

Suppose that a sequence has a fraction of sites *m_1_* evolving at rate *μ*_1_ = *μ* and the remaining sites (1 − *m*_1_) evolving epistatically such that a pair of sites need to mutate simultaneously for the offspring to recover wild-type fitness, i.e. *μ*_2_ = *μ*^2^. Assuming the saturation frequency across all sites are equal and that the model correctly identifies their frequency, i.e. *α_i_* = *α* = *α_M_*, we can use **Equation S2** to recover the expected substitution rate 〈*μ*〉 = *m*_1_ *μ* + (1 - *m*_1_)*μ*^2^. As the fast-evolving sites approach the saturation point at *t*_1_ ≈ *α*/*μ*, the inflection point of rate decay emerges and a sharp decline in estimated substitution rate follows while the remaining fraction of sites, (1 − *m*_1_), keep accumulating new substitutions at rate *μ*^2^, slowing down the slope of rate decay until those sites also reach saturation at *t*2 ≈ *α*/*μ*^2^ beyond which point the entire genome reaches saturation and the power-law rate decay with slope −1 emerges. We can also see from **Figure S1 b** that as the proportion of slow-evolving sites increases, the mean substitution rate goes down and the slope of the time-dependent rate decay becomes less steep.

### Simulating a Wright-Fisher population to infer substitution rates

We simulate a neutral haploid Wright-Fisher population of size *N_e_* with *L* evolving sites under a constant mutation rate *μ* per site such that every nucleotide (A, C, G, and T) can mutate to any other nucleotide at the same rate *μ*/3 – mutation rate is equal to substitution rate under neutrality. We then sample from the entire population at two time points with an increasingly wider time gap, *t*\*. Initially, we allow the population to evolve for 10*N_e_* generations before taking the first sample to ensure that neutral coalescent events reach their steady state distribution and that the population, on average, coalesce every 2*N_e_* generations. We then take the second sample *t*\* generations later and repeat this process 100 times to generate replicate sequences at both time points and run each set of simulations in BEAST 1.10 to estimate the substitution rate (**Figure S2**). We load the simulated sequences (along with their sampling times) on BEAST and use a strict molecular clock with a continuous-time Markov chain reference prior on substitution rates, a constant population coalescent prior, and a Jukes-Cantor substitution model. For every simulated set, the Markov chain Monte Carlo was run for 10,000,000 steps and parameter convergence was inspected visually.

In **Figure S2**, we recreate the time-dependent pattern of rate decay both in the absence and presence of rate heterogeneity across sites, using a standard substitution model on simulated data. We find that while the inferred substitution rates exhibit a power-law rate decay with slope −1 over longer time intervals (see **Figure S2 (a)** and **(b)**), the inferred TMRCAs tend to be overestimated with a similar (inverse) power-law trend, i.e. 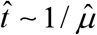(see **Figure S2 (c)** and **(d)**). We also find an unexpected time-dependent rate effect over short timescales. This occurs when the observation gap, *t*\*, is much shorter than the expected coalescent time of the population, i.e. *t*\* ≪ 2*N_e_*. This also results in the underestimation of true TMRCAs which systematically makes worse predictions for higher substitution rates. The expected rate curves (dashed lines shown in **Figure S2 (a)** and **(b)**) can be approximated by replacing *p*(*t*) from **Equation S1** into 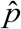 from **Equation S2** which is given by

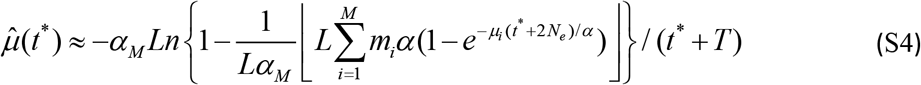

Such ⌊·⌋ is the floor function which represents the finite size effect of having *L* evolving sites on saturation frequency. The mean divergence time between the two populations is approximately *t* ≈ *t*\* + 2*N_e_* and the inferred divergence time is 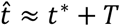 – this resembles the mis-calibration effects reported elsewhere (see Equation 2 in ref^55^). The reason why **Equation S4** only works as an approximate is that the median inferred TMRCA from simulation results, *T*, also varies with respect to observation gap *t*\* (see **Figure S2 (c)** and **(d)**). However, for *t*\* ≫ 2*N_e_*, the variation in *T* becomes negligible compared to *t*\*and only has second-order effects on inferred substitution rates. **Figure S2 (e)** and **(f)** show the tree topology under the two extremes, *t*\* ≪ 2*N_e_* and *t*\* ≫ 2*N_e_*, respectively. It indicates that, over long timescales, the time-dependent rate effects are dominated by the very long (and saturated) branch connecting the two populations that are *t*\* generations apart. As a result, the decay dynamics looks very similar to the analytical results in **Figure S1** where we estimate the substitution rate between a pair of sequences separated by a very long branch.

#### Supplementary Figures

**Figure S1:**
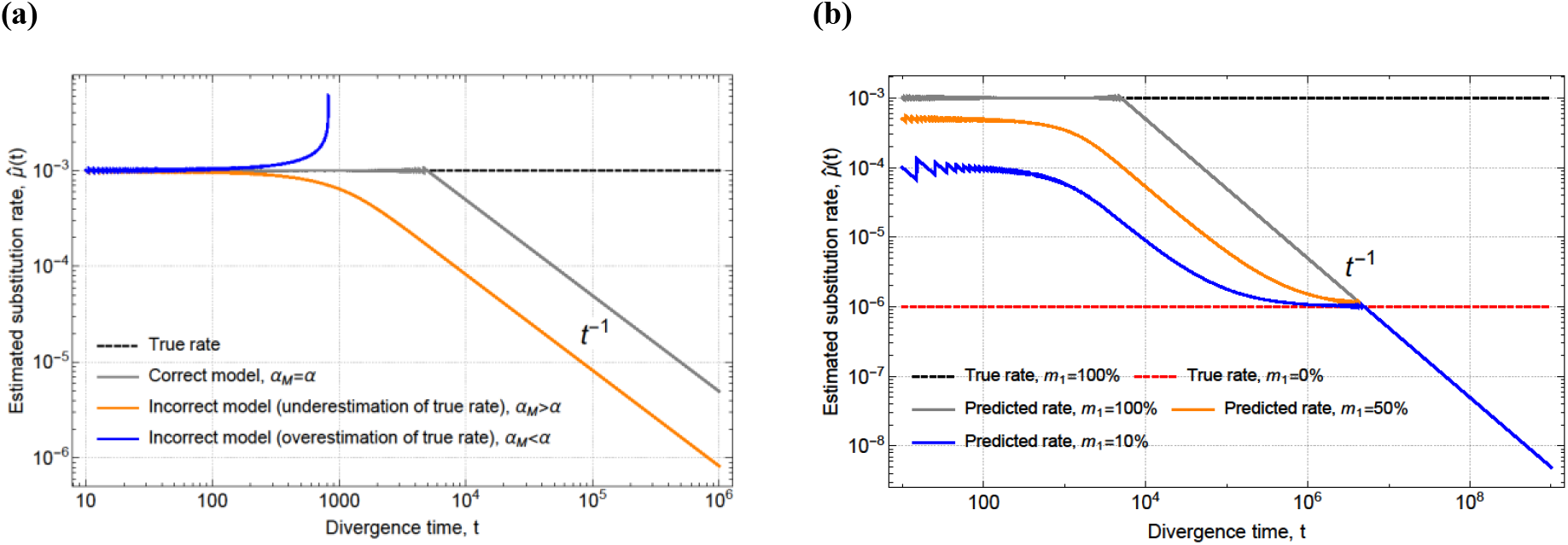
**(a)** Estimating the substitution rate *μ* = 10^−3^ (black dashed line) using **Equation S2** for a pair of sequences that have diverged from each other *t* generations ago: using the correct substitution model (gray), a model that over-estimates the true saturation frequency, *α*, (orange), and a model that underestimates the true saturation frequency (blue). **(b)** Estimating the expected (or mean) substitution rate, < *μ* >, when a fraction *m*_1_ of sites evolve at rate *μ*_1_ = 10^−3^ (black dashed line) and the remaining fraction, 1 − *m*_1_, at rate *μ*_2_ = 10^−6^ (red dashed line). The expression *t*^−1^ in both plots shows the dominating term in rate decay with respect to divergence time, *t*, corresponding to slope −1 on the graphs.

**Figure S2:**
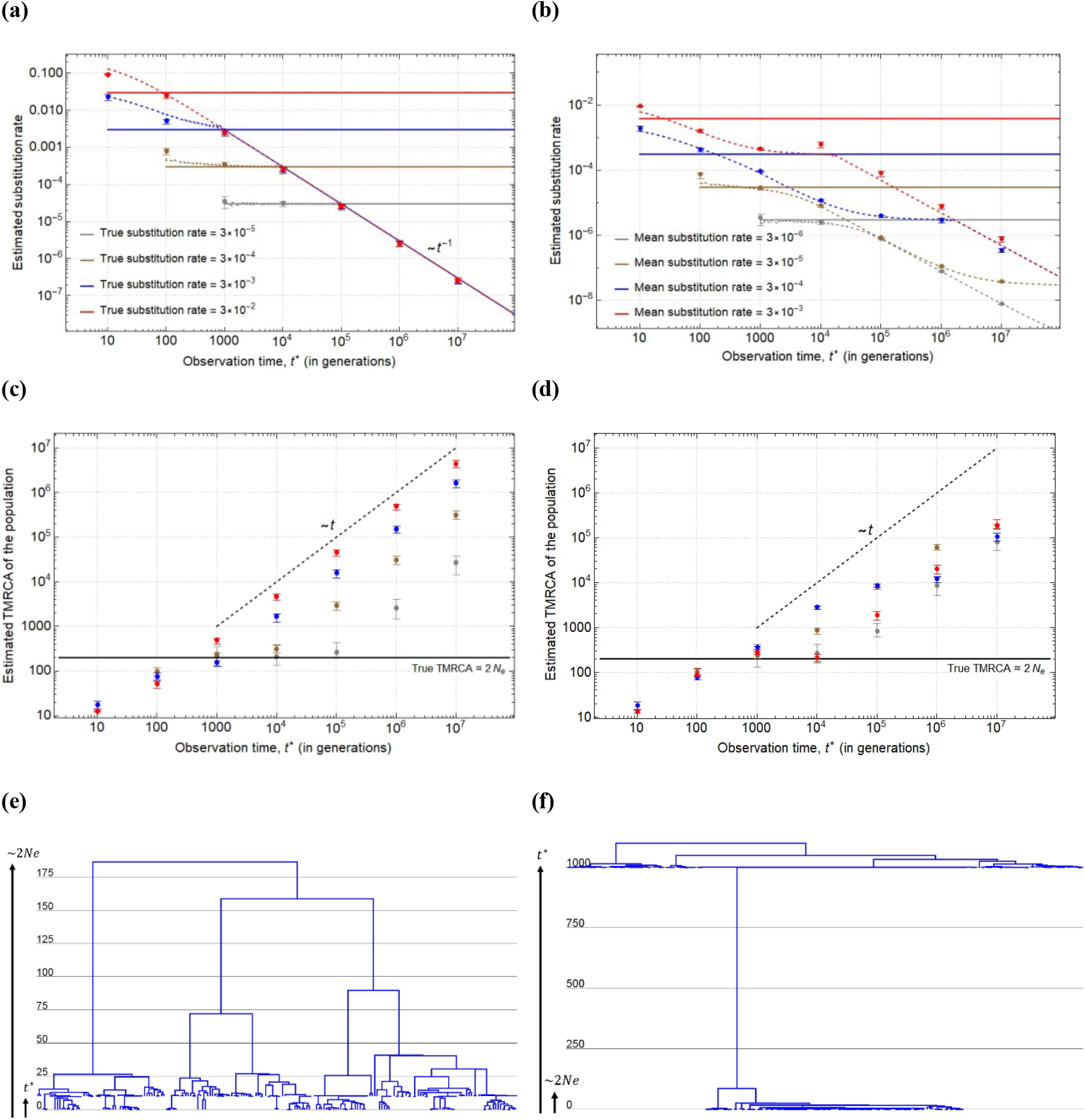
Estimated substitution rate for (**a**) a population of size *N_e_* evolving according to a neutral model of substitution where *L_n_* = 100 sites evolve at the same rate, *μ*, and (**b**) a model with rate heterogeneity across sites such that *L*_1_ = 100 sites evolve at rate *μ* and the remaining *L*_2_ = 900 sites at rate *μ*^2^ measured as a function of observation gap, *t*\*, between when the first and second sample is taken from the population. Dashed lines show the theoretical prediction according to **Equation S4** and solid lines show the mean (expected) rates used for the simulations. (**c**) and (**d**) show the estimated TMRCA for the first group of sampled sequences, i.e. at *t*\* = 0, and solid lines show the mean TMRCA according to neutral theory. The rates and TMRCAs are estimated using BEAST 1.10 under a strict clock assumption (see Methods section). Dots represent the median value taken from 100 independent runs and error bars show the interquartile region. The maximum clade credibility trees for one simulation run corresponding to *μ* = 3 × 10^−5^ is shown when the observation gap is (**e**) *t*\* = 10, *t*\* ≪ 2*N_e_* and (**f**) *t*\* = 1000, *t*\* ≫ 2*N_e_*.

**Figure S3:**
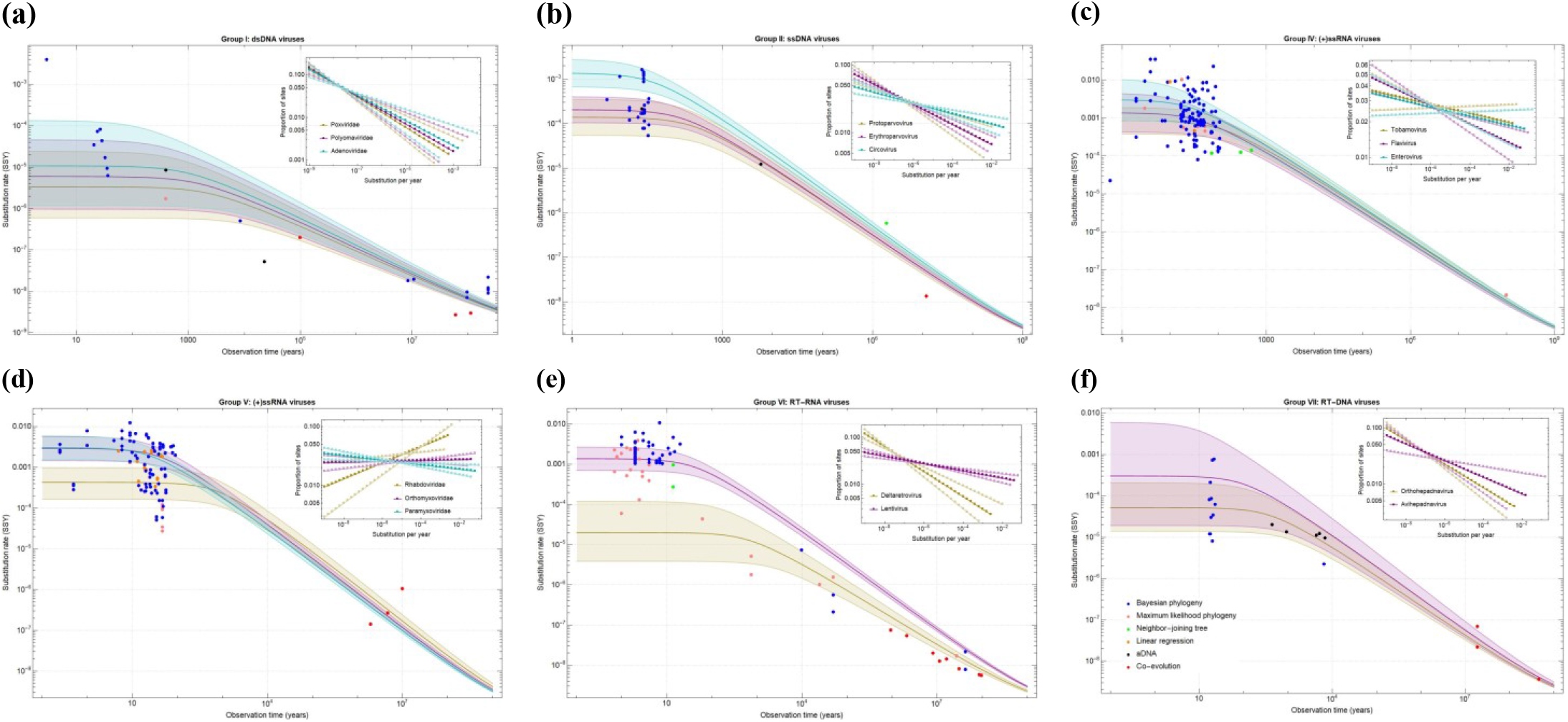
(**a**)-(**f**) Estimated time-dependent rate curves for each viral group according to the PoW model and their corresponding distribution of rate groups (inset). Two or three distinct mean substitution rates (coloured in gold, purple, and blue) are selected for each virus family to estimate rate curves according to the PoW model. The solid lines show the best fit and shaded areas the 95% confidence interval ( Δ*M* = 1.58 and *α_M_* = *α* = 3 / 4).

**Figure S4:**
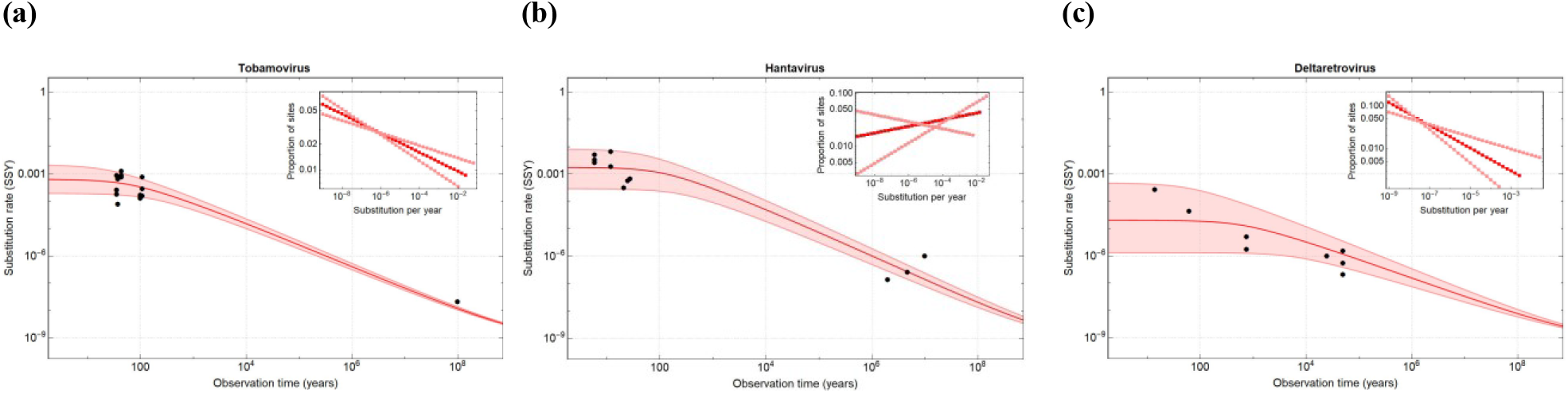
Estimated time-dependent rate curves for three selected genera. (**a**) Tobamoviruses with 〈*μ*〉 = 0.6(0.2 - 2) ×10^−3^ SSY and *μ*_max_ = 3(1 - 6) ×10^−2^ SSY, (**b**) Hantaviruses with 〈*μ*〉 = 2(0.3 - 8) ×10^−3^ SSY and *μ*_max_ = 2(0.6 - 4) ×10^−2^ SSY, and (**c**) Deltaretroviruses with 〈*μ*〉 = 2(0.1 - 50) ×10^−5^ SSY and *μ*_max_ = 3(2 - 30) ×10^−3^ SSY. These genera are selected as they have the largest timespan of rate measurements.

**Table S1:** Estimated mean and maximum substitution rate according to the PoW model using different virus families or genera to calibrate the mean rate across 6 viral groups. Parentheses correspond to 95% confidence intervals.

| Viral group | Virus family/genus used for calibration* | Mean substitution rate, $\langle \mu \rangle$ | Fastest rate group, $\mu_{\max}$ |
| --- | --- | --- | --- |
| Group I | Poxviridae | $0.3(0.06 - 2) \times 10^{-5}$ | $0.6(0.2 - 3) \times 10^{-3}$ |
| | Polyomaviridae | $0.6(0.1 - 5) \times 10^{-5}$ | $1(0.3 - 4) \times 10^{-3}$ |
| | Adenoviridae | $1(0.1 - 10) \times 10^{-5}$ | $2(0.3 - 10) \times 10^{-3}$ |
| Group II | Protoparvovirus | $1(0.6 - 4) \times 10^{-4}$ | $0.6(0.4 - 1) \times 10^{-2}$ |
| | Erythroparvovirus | $2(1 - 4) \times 10^{-4}$ | $1(0.6 - 2) \times 10^{-2}$ |
| | Circovirus | $10(7 - 300) \times 10^{-4}$ | $4(3 - 6) \times 10^{-2}$ |
| Group IV | Tobamovirus | $0.4(0.9 - 2) \times 10^{-3}$ | $2(1 - 3) \times 10^{-2}$ |
| | Flavivirus | $1(0.4 - 5) \times 10^{-3}$ | $4(2 - 10) \times 10^{-2}$ |
| | Enterovirus | $3(0.9 - 10) \times 10^{-3}$ | $6(3 - 10) \times 10^{-2}$ |
| Group V | Rhabdoviridae | $0.4(0.2 - 1) \times 10^{-3}$ | $0.3(0.2 - 0.4) \times 10^{-2}$ |
| | Orthomyxoviridae | $3(1 - 6) \times 10^{-3}$ | $4(3 - 6) \times 10^{-2}$ |
| | Paramyxoviridae | $3(2 - 6) \times 10^{-3}$ | $6(4 - 10) \times 10^{-2}$ |
| Group VI | Deltaretrovirus | $0.02(0.004 - 0.1) \times 10^{-3}$ | $0.3(0.06 - 1) \times 10^{-2}$ |
| | Lentivirus | $1(0.7 - 3) \times 10^{-3}$ | $4(3 - 6) \times 10^{-2}$ |
| Group VII | Orthohepadnavirus | $5(1 - 20) \times 10^{-5}$ | $4(2 - 10) \times 10^{-3}$ |
| | Avihepadnavirus | $30(2 - 600) \times 10^{-5}$ | $20(2 - 100) \times 10^{-3}$ |

\*Only short-term rate estimates (measured over time scales of <100 years) – along with the long-term rate estimates (>100 years) from the entire data set – from this particular virus family or genus (rather than the entire viral group) is used for the calibration.

